# Multifaceted effects on *even-skipped* transcriptional dynamics upon *Krüppel* dosage changes

**DOI:** 10.1101/2023.06.27.546766

**Authors:** Shufan Lin, Bomyi Lim

## Abstract

Although fluctuations in transcription factor (TF) dosage are often well tolerated, little is known about how TF dosage modulation changes the target gene expression dynamics and results in subsequent developmental phenotypes. Using MS2/MCP-mediated quantitative live imaging in early *Drosophila* embryos, we characterized transcriptional dynamics of the pair-rule gene *even-skipped* (*eve*) upon changing the gap gene *Krüppel* (*Kr*) level. Decreasing *Kr* dosage by half leads to a transient posterior expansion of the *eve* stripe 2 and an anterior shift of stripe 5. Surprisingly, the most significant changes are observed in *eve* stripes 3 and 4, whose enhancers do not contain Kr binding sites. For both stripes, the widths of stripes are narrowed, boundaries are located more anteriorly, and the mRNA production level is reduced in *Kr* heterozygous embryos. We show that Kr dosage indirectly affects stripes 3 and 4 dynamics by modulating other gap gene dynamics. We quantitatively correlate moderate body segment phenotypes of *Kr* heterozygotes with spatiotemporal changes in *eve* expression. Our results indicate that nonlinear relationships between TF dosage and phenotypes underlie direct TF-DNA and indirect TF-TF interactions.

**Summary statement:** This study analyzed the changes in transcriptional dynamics of *even-skipped* upon halving *Krüppel* dosage, and revealed how the combination of direct TF-DNA and indirect TF-TF interactions shape proper patterning.

## Introduction

Body segmentation is a critical process that specifies repeated units along the anterior-posterior (AP) axis during early embryogenesis. Although detailed processes differ among species, body segmentation is a conserved strategy to create blueprints to guide the formation of structures and organs (Akam, 1987; Diaz-Cuadros et al., 2021; Saga and Takeda, 2001). In *Drosophila*, body segment formation is completed through a hierarchical gene regulatory network that encodes cascades of transcription factors (TFs) (Ingham, 1988; Jaeger, 2011; Schroeder et al., 2011). *bicoid* (*bcd*), *nanos*, and other maternal effect genes are first localized to either the anterior or posterior ends and polarize the embryo, establishing sequential gap gene domains in a concentration-dependent manner along the AP axis (Driever and Nüsslein-Volhard, 1988; Ephrussi and Johnston, 2004; Rivera-Pomar et al., 1995; Wang et al., 1994). Refined expression of gap genes gives rise to distinct stripes of pair-rule genes in nuclear cycle 14 (NC14) (Goto et al., 1989; Surkova et al., 2008). In later stages, pair-rule genes regulate segment-polarity genes and complete the body segmentation process (DiNardo and O’Farrell, 1987; French, 2001). The expression of downstream patterning genes is tightly regulated by the spatiotemporal dynamics of upstream gene products. For example, activation and repression by multiple gap genes cause a pair-rule gene *even-skipped* (*eve*) to be expressed in seven stripes, where each stripe is regulated by a stripe-specific enhancer containing different combinations of gap gene TF binding sites (Fujioka et al., 1999; Harding et al., 1989; Small et al., 1992; Small et al., 1996).

Yet, *Drosophila* often exhibits high tolerance to TF dosage perturbations. Flies carrying three or four copies of *bcd* can buffer the high dosage through the interplay among downstream gap genes and develop into normal adults (Berleth et al., 1988; Liu et al., 2013). In addition, heterozygous mutants for most patterning genes are viable and fertile (Driever and Nüsslein-Volhard, 1988; Surkova et al., 2019; Yu and Small, 2008). On the other hand, such high tolerance does not necessarily imply robustness of embryonic development. Some larvae that are heterozygous for *hunchback* (*hb)* or *Kr* were found to have partial body segment deletions (Lehmann and Nüsslein-Volhard, 1987; Wieschaus et al., 1984). Phenotypic differences due to moderate changes in TF dosage were also observed in other species. In humans and mice, reduced dosage of a TF *Sox9* drives skeletal malformations at the lower jaw (Long et al., 2020; Naqvi et al., 2023). These results indicate that some degree of robustness to different TF dosages are conserved across species, preventing severe phenotypic consequences like lethality; yet, expression of downstream genes can still be altered upon TF dosage modulations to induce moderate non-lethal phenotypes.

We use the highly dynamic interactions among gap genes and pair-rule genes to study the role of TF dosage in patterning and developmental robustness. Both gap and pair-rule genes undergo stochastic transcription at the beginning of NC14, followed by the formation and refinement of sharp expression domains in 50 minutes (Jaeger et al., 2004; Perry et al., 2012; Surkova et al., 2008). Although flies with reduced gap gene copies are viable, little is known about how the change in TF dosage affects the dynamics of pair-rule gene regulation and subsequent developmental robustness. Does a decrease in TF concentration drive ectopic target gene expression? Are these changes connected to the defects observed in some heterozygotes from previous studies? With a real-time, high spatiotemporal resolution platform, we can quantify subtle changes in transcriptional activity to elucidate the molecular mechanism of how fluctuations in TF dosage lead to phenotypic consequences.

In this study, we utilized MS2/MCP and PP7/PCP-based live imaging and genetic perturbations to examine the direct and indirect role of a gap gene *Kr* dosage in the transcriptional regulation of a pair-rule gene *eve*. We quantified distinct changes of different *eve* stripes in response to the decrease in *Kr* dosage. The observed shift in *eve* stripe 2 and 5 boundaries are presumably due to the reduced Kr binding to the respective enhancers. Surprisingly, the most affected domains are *eve* stripes 3 and 4, even though their enhancers lack Kr binding sites. Decreased Kr shifts stripe boundaries anteriorly, narrows the expression domain, and reduces mRNA production. We further demonstrate that these effects on stripes 3 and 4 are propagated by the changes of other gap genes in *Kr* heterozygotes. Halved *Kr* dosage induces more anteriorly located giant (*gt*) and *knirps* (*kni*) expression, and a significant drop in *hb* expression level compared to wild-types. Since stripes 3 and 4 are directly regulated by Hb and Kni, we propose that Kr indirectly affects stripes 3 and 4 by altering the dynamics of adjacent gap gene domains. By quantitatively analyzing how individual nuclei respond to fluctuations in Kr concentration, our results provide insights into the systematic interplay among multiple genes in determining body pattern formation.

## Results

### Decreased *Kr* directly affects the formation of *eve* stripes 2 and 5

We utilized a dual-color MS2 and PP7 system to visualize endogenous *eve* transcription activity in wild-type and *Kr* heterozygous (*Kr^1^/+*) *Drosophila* embryos (Fig. 1A, Fig. S1A-B, Movie S1) (Garcia et al., 2013; Hocine et al., 2013; Lim et al., 2018a). Heterozygotes and wild-types can be distinguished with the absence and presence of the *iab5>PP7* reporter gene expression, respectively (Fig. 1A). While all seven *eve* stripes are formed by late NC14 for *Kr* heterozygous embryos, both spatial and temporal dynamics of the stripe formation (Fig. 1B, Fig. S1D-E), and the subsequent mRNA production level (Fig. 1C, Fig. S1C) are affected.

**Fig. 1.**
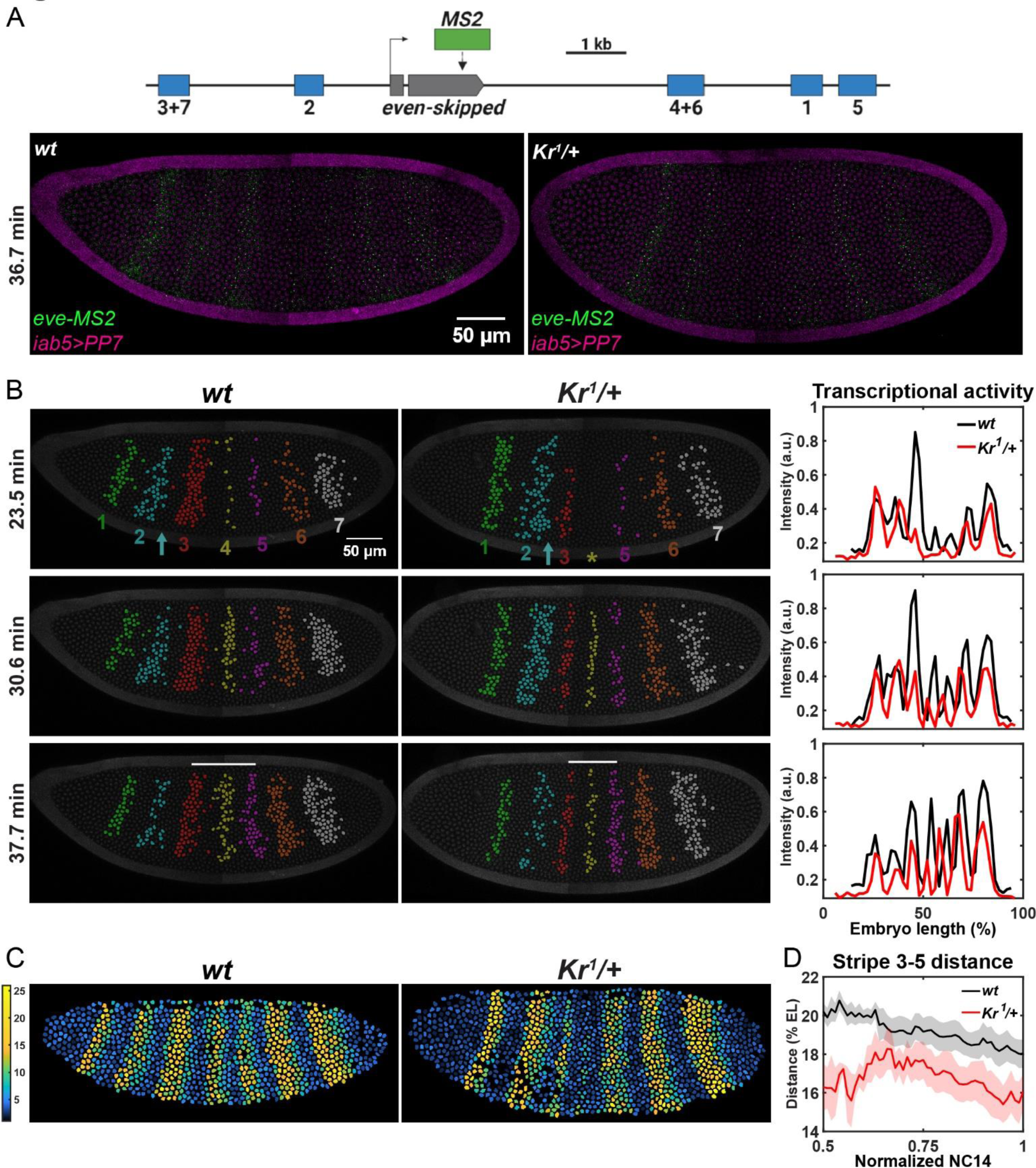
Halved *Kr* dosage affects the overall transcriptional dynamics of *eve*. (A) Top: schematic of the *eve-MS2* construct. Bottom: Snapshots of a wild-type and *Kr* heterozygous embryo expressing *eve-MS2* (green) and *iab5>PP7* (magenta) in NC14. (B) Snapshots of a wild-type and *Kr* heterozygous embryo expressing *eve-MS2*. Transcriptionally active nuclei are false-colored. Nuclei are labeled with His2Av-eBFP2 (dark gray). Yellow star - position of *eve* stripe 4; cyan arrows - region between stripes 2 and 3; white bars - distance between stripes 3 and 5. Right panel shows transcriptional activity of *eve-MS2* across the AP axis from the left panels. (C) Heatmaps showing cumulative mRNA production in individual nuclei. (D) Average distance between *eve* stripes 3 and 5 over time. Error bar represents mean ± s.e.m. of wild-type (n=7) and *Kr* heterozygous (n=5) embryos.

To better quantify the effects of *Kr* dosage on *eve*, we focused on the transcriptional activity of stripes 2-5, which overlap with the Kr expression domain. Transcription of the seven *eve* stripes is regulated by five stripe-specific enhancers, and Kr directly binds to the stripe 2 and 5 enhancers (Fig. 1A) (Fujioka et al., 1999; Harding et al., 1989; Small et al., 1992; Small et al., 1996). As a result, reduced *Kr* dosage affects the spatial pattern of these two stripes. In mid NC14, stripe 2 shows posterior expansion, such that the width of stripe 2 is ∼1.6 nuclei wider in *Kr* heterozygotes (Fig. 1B, cyan arrow). However, this expansion is only transient, as the stripe width becomes comparable between wild-types and heterozygotes in late NC14 (Fig. 2A-C). We hypothesize that the reduced Kr concentration induces stripe 2 to expand in mid NC14 (Fig. 2A, C). However, the anterior shift of *Kr* domain and the increasing level of Kr protein during NC14 (Crombach et al., 2012; Surkova et al., 2008) allow expanded *eve* stripe 2 domain in heterozygotes to be refined to match the wildtype width in late NC14 (Fig. 2B-C). Similarly, *eve* stripe 5 domain is slightly wider in *Kr* heterozygous embryos, despite more embryo-to-embryo variability (Fig. 2A-B, D). Due to the transient domain expansion (stripe 2) and high variability in width (stripe 5), the total number of transcriptionally active nuclei in stripe 2 and 5 domains remain comparable between wild-types and heterozygotes (Fig S2 A-D). Reduced Kr concentration also shifted the position of stripe 5. While stripe 2 domain does not change significantly, stripe 5 is located ∼3 nuclei closer to the anterior tip in *Kr* heterozygotes than in wild-types (Fig. 2E-F). Despite changes in boundary positions, the amount of mRNA produced by individual nuclei, their transcriptional amplitude, and duration of active transcription in these two stripes are similar between wild-type and heterozygous embryos (Fig. 1C, Fig. 2G-J, Fig. S2E-F). In summary, our results indicate that stripe 2 and 5 enhancers are sensitive to the decrease in Kr concentration, but the impact of Kr is limited to shifts in boundary positions, with little effect on the mRNA production of both stripes.

**Fig. 2.**
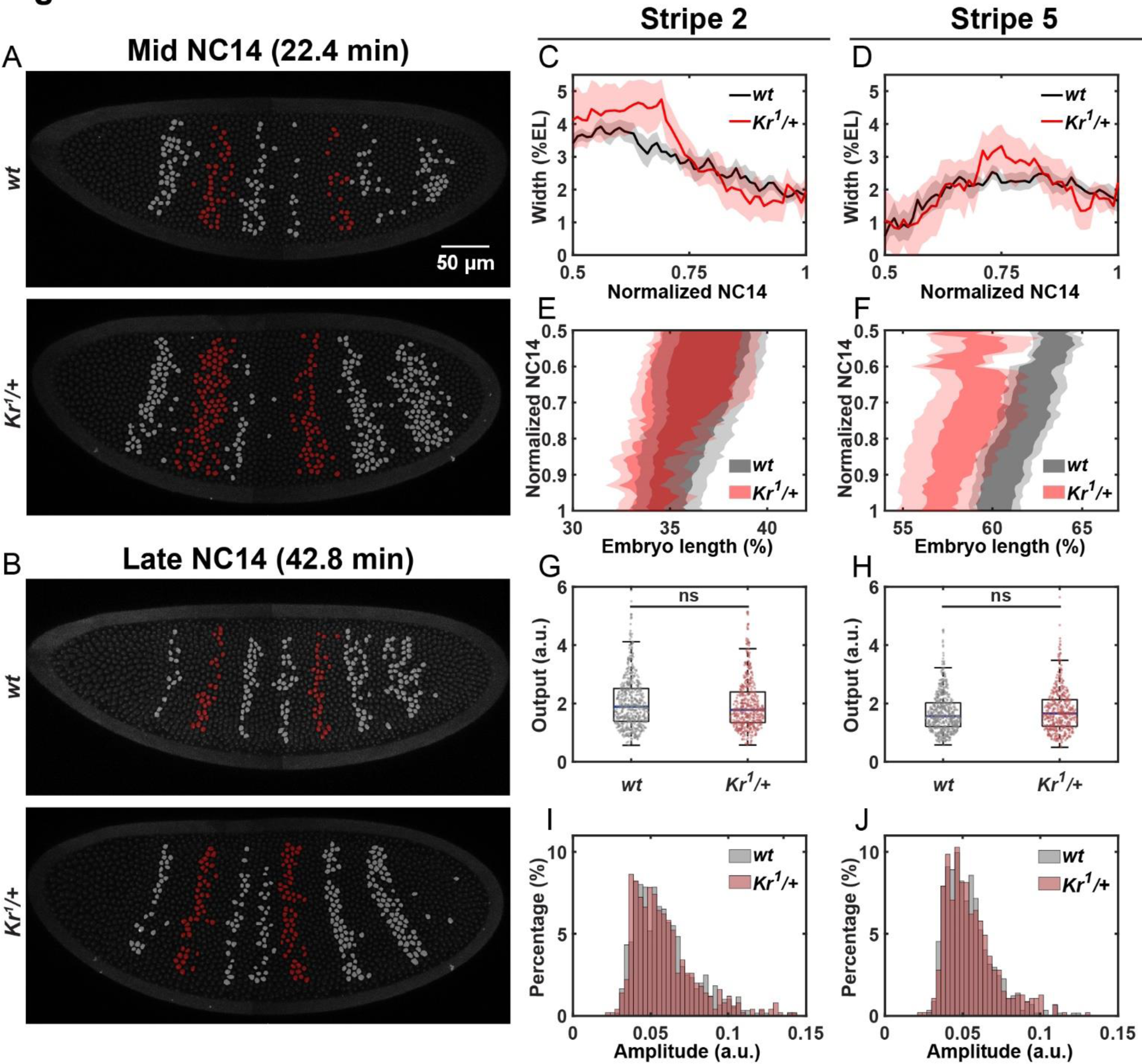
Decreased *Kr* dosage directly changes *eve* stripes 2 and 5 boundary positions. (A-B) False-colored wild-type and *Kr* heterozygous embryos. Transcriptionally active nuclei within the *eve* stripe 2 and 5 domains are in red, and other active nuclei are in gray. (C-D) Average width of stripe 2 (C) and 5 (D) over time. (E-F) Average position of stripe 2 (E) and 5 (F). Error bars represent mean ± s.e.m. for (C)-(F). (G-H) Cumulative mRNA output of individual nuclei within stripe 2 (G) and 5 (H). Scatters represent individual nuclei. ns, not significant from the Student’s t-test. (I-J) Distribution of the average transcriptional amplitude of individual nuclei in stripe 2 (I) and 5 (J). 599 (stripe 2 *wt*), 498 (stripe 2 *Kr^1^/+*), 572 (stripe 5 *wt*), and 467 (stripe 5 *Kr^1^/+*) from 7 wild-type and 5 *Kr* heterozygous embryos were analyzed.

### Decreased *Kr* indirectly hampers *eve* transcription in stripe 3 and 4 domains

We observed the most significant changes in stripes 3 and 4 (Fig. 1B and Fig. 3A). Stripe 4 formation is delayed by ∼6 min, and much fewer nuclei comprise the stripe (Fig. 1B - yellow star, Fig. 3C, Fig. S3B). Stripe 3 has a similar timing of activation in both wild-types and heterozygotes, but fewer nuclei are activated in heterozygotes’ domain (Fig. 3A-B, Fig. S3A). As a result, the width of stripes 3 and 4 are both narrower in heterozygotes (Fig. 3D-E). We note that the nuclei located in the posterior region of stripe 3 are not activated in heterozygotes, thus affecting the posterior boundary of stripe 3 (Fig. 3F). For stripe 4, the entire expression domain shifts anteriorly in heterozygotes compared to wild-types (Fig. 3G). Combined with an anteriorly located stripe 5 (Fig. 2F), the distance from stripe 3 to 5 is narrower in *Kr* heterozygous embryos (Fig. 1B - white bar, Fig. 1D), agreeing with previous studies (Surkova et al., 2013).

**Fig. 3.**
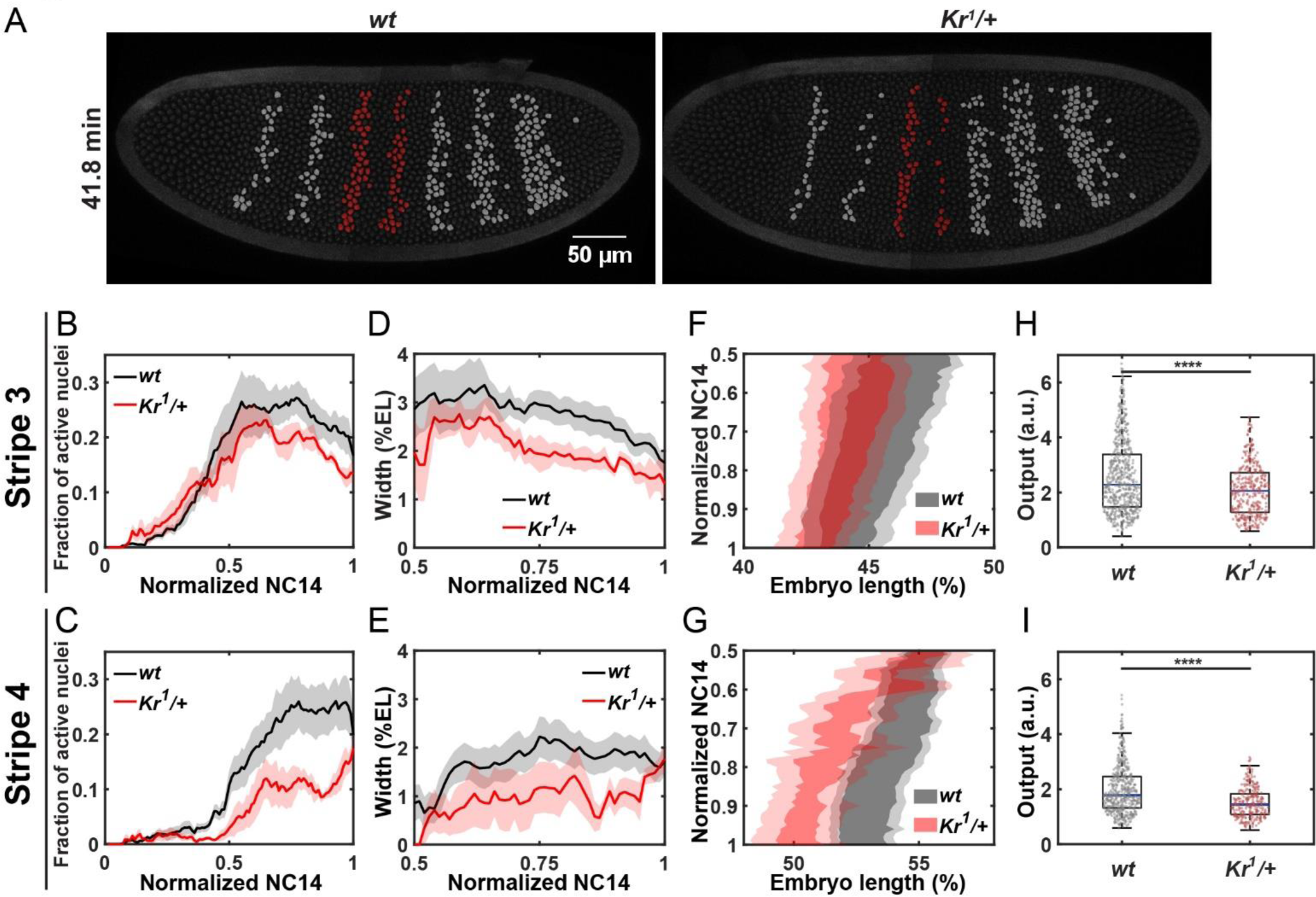
Decreased *Kr* dosage affects *eve* stripes 3 and 4 boundary positions and transcriptional activity. (A) False-colored wild-type and *Kr* heterozygous embryos. Transcriptionally active nuclei within the *eve* stripe 3 and 4 domains are in red, and other active nuclei are in gray. (B-C) Average activation kinetics of stripe 3 (B) and 4 (C). (D-E) Average width of stripe 3 (D) and stripe 4 (E). (F-G) Average position of stripe 3 (F) and stripe 4 (G). Error bars represent mean ± s.e.m.. (H-I) Cumulative mRNA output of individual nuclei within stripe 3 (H) and stripe 4 (I). Scatters represent individual nuclei. ****P<0.0001 from the Student’s t-test. 617 (stripe 3 *wt*), 377 (stripe 3 *Kr^1^/+*), 488 (stripe 4 *wt*), and 281 (stripe 4 *Kr^1^/+*) from 7 wild-type and 5 *Kr* heterozygous embryos are analyzed.

In addition to changes in spatial dynamics, fewer mRNA are accumulated in stripe 3 and 4 regions in heterozygotes than in wild-types (Fig. 1C, Fig. 3H-I). Further investigation suggests that this decrease is a result of both shortened active transcriptional duration and lower transcriptional amplitude, although the contribution of duration is moderate for stripe 3 (Fig. S3C-F). Moreover, we observed sporadic transcriptional activity of stripe 4 nuclei in *Kr* heterozygotes (Movie S2). This sporadic expression not only contributes to the most disrupted expression in stripe 4 compared to other affected stripes (Fig. 1C, Fig. S1C), but also causes the high variability in stripe 4’s spatial pattern (Fig. 3G).

A previous study found high penetrance of denticle band defects in mesothorax, metathorax, and the second abdominal segment of *Kr* heterozygous larvae (Wieschaus et al., 1984). Indeed, the expression of *eve* stripes 3 and 4 are involved in defining these segments (Lawrence et al., 1987; Martinez-Arias and Lawrence, 1985). Our quantification of *eve* transcriptional profiles implies that narrowed stripe width and reduced mRNA production of these two stripes fail to provide precise positional information for the downstream genes to complete proper body segmentation in *Kr* heterozygous embryos.

### Decreased *Kr* regulates *eve* stripes through adjacent gap genes

ChIP-ChIP and ChIP-seq analyses show strong Kr binding at the *eve* 2 and 5 enhancers, indicating direct impact of Kr on *eve* stripes 2 and 5. However, there is no Kr binding at the 4+6 enhancer and very low Kr binding at the *eve* 3+7 enhancer (Castro-Mondragon et al., 2022; Li et al., 2008; Paris et al., 2013). Since stripe 3 expression can be reconstructed without Kr *in silico* (Ilsley et al., 2013), we assumed that *eve* stripes 3 and 4 are not directly regulated by Kr. Instead, ChIP-seq and misexpression analyses provided evidence of Hb and Kni directly repressing stripes 3 and 4 (Clyde et al., 2003; Paris et al., 2013). As mutual repression exists among Kr, Kni, and Hb, the anterior shift of stripes 3 and 4 boundaries in *Kr* heterozygotes compared to wild-types may be caused by an indirect effect of decreased *Kr* influencing its adjacent *kni* and *hb* domains (Capovilla et al., 1992; Clyde et al., 2003; Hülskamp et al., 1990; Sauer and Jäckle, 1991).

To further investigate this hypothesis, we used RNA fluorescence *in situ* hybridization (FISH) and live imaging to quantify the expression of *hb*, *kni*, and *gt*, three gap genes that regulate the transcription of *eve* stripes 2-5. For all the gap genes analyzed, we assumed a linear correlation between mRNA and subsequent protein translation (Becker et al., 2013; Bothma et al., 2018). Since both maternal and zygotic Hb regulate downstream genes, we performed FISH to visualize total *hb* mRNAs. A low level of Kr was shown to activate *hb* expression (rather than working as a repressor) (Holloway and Spirov, 2015; Sauer and Jäckle, 1991). Indeed, we observed decreased *hb* concentration across the AP axis at mid NC14 with no significant changes in expression domains (Fig. 4A, D). Since Hb proteins bind to the 4+6 enhancer as repressors, the decreased *hb* expression in *Kr* heterozygotes may cause the anterior shift of the stripe 4 (Fig. 4G, purple) (Clyde et al., 2003).

**Fig. 4.**
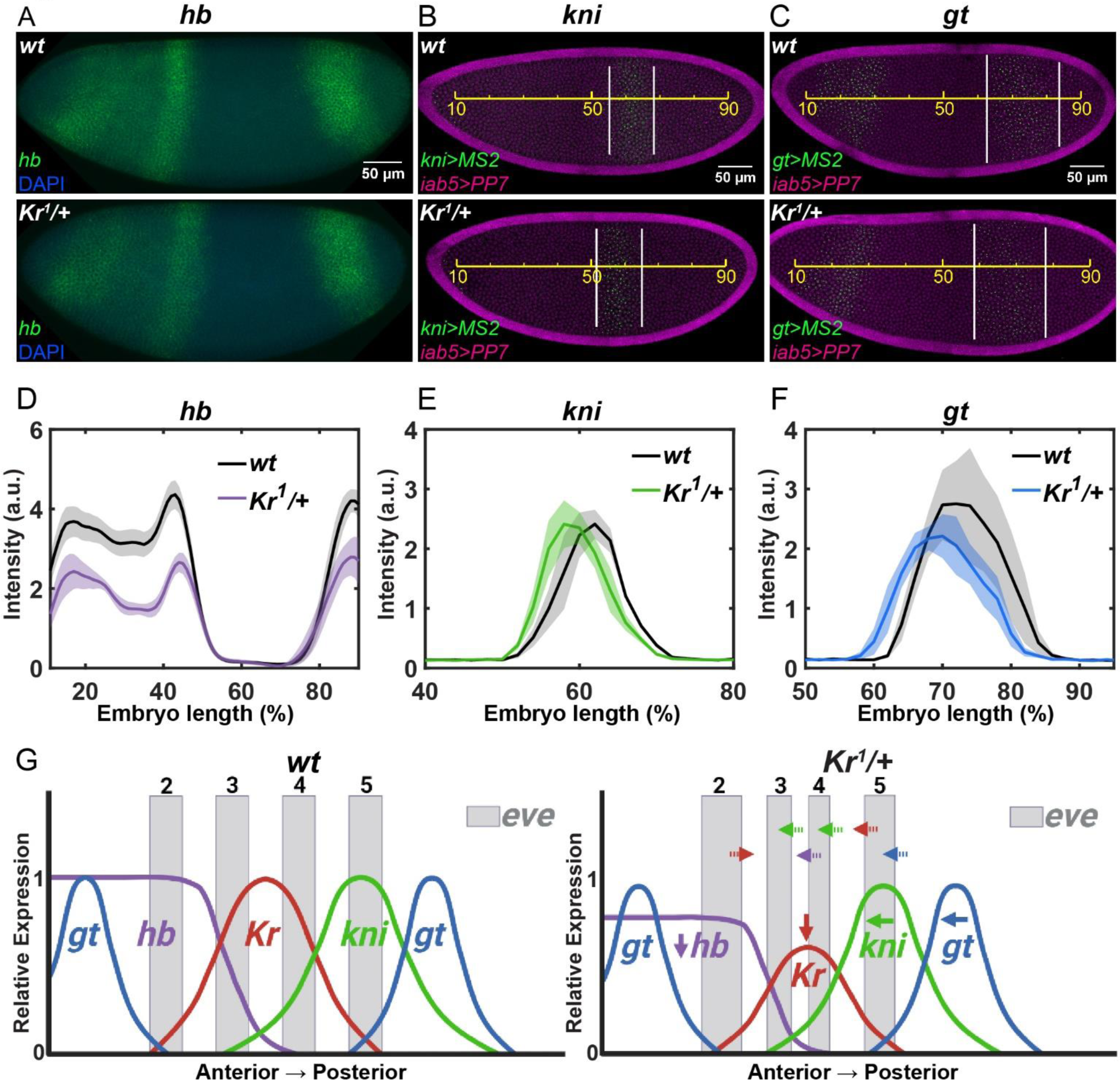
Decreased *Kr* dosage affects adjacent gap genes. (A) Wild-type and *Kr* heterozygous embryos stained with *hb* (green) and DAPI (blue). (B-C) A wild-type and *Kr* heterozygous embryos expressing *kni>MS2* (green) and *iab5>PP7* (magenta) (B) and *gt>MS2* (green) and *iab5>PP7* (magenta) (C). Yellow line - EL; white line - *kni>MS2* or *gt>MS2* expression domain. (D-F) Spatial profile of *hb* (D), *kni* (E), and *gt* (F) transcriptional activity. Error bars show the mean ± s.e.m.. 10, 3, and 3 wild-type and 7, 3, and 3 *Kr* heterozygous embryos were used to visualize *hb, kni>MS2,* and *gt>MS2*, respectively. (G) A model illustrating how the halved *Kr* dosage affects gap genes and *eve* expressions. Decreased *Kr* shifts the posterior and anterior boundary of *eve* stripes 2 and 5, respectively (red). Decreased *hb* in *Kr* heterozygotes induces anterior shift of the *eve* stripe 4 (purple). Anterior shift of *kni* and *gt* induces the anterior shift of stripe 3 and 4 posterior boundaries (green) and stripe 5 posterior boundary (blue).

Since Kni directly defines the posterior boundaries of stripes 3 and 4 by binding to *eve* 3+7 and 4+6 enhancers, we next examined changes in *kni* expression (Clyde et al., 2003). Spatial profiles of *kni* in *Kr* heterozygotes reveal a more anteriorly located domain compared to the wild-types (Fig. 4B, E). This pushes the posterior boundaries of stripes 3 and 4 to the anterior side in *Kr* heterozygous embryos (Fig. 4G, green). In mid-NC14, a high level of *kni* is expressed more anteriorly in heterozygotes, repressing the transcription of stripe 4. In late NC14, however, *eve* autoregulatory enhancer regulates the late activation in stripe 4 nuclei, and restores stripe 4 expression towards the end of NC14 (Fig. 3C) (Goto et al., 1989; Jiang et al., 1991).

Lastly, we examined the changes in *gt* expression dynamics upon *Kr* dosage modulation. As expected, the posterior *gt* stripe is expressed more anteriorly in the heterozygous embryos (Fig. 4C, F), while the anterior stripes remain unchanged (Fig. S4) due to the lack of Kr binding sites in the gt-23 enhancer that regulates anterior *gt* stripes (Hoermann et al., 2016; Ochoa-Espinosa et al., 2005). Gt directly binds to *eve* stripe 2 and 5 enhancers to regulate the anterior boundary of stripe 2 and the posterior boundary of stripe 5 (Fujioka et al., 1999; Small et al., 1991). Therefore, the anteriorly expressed *gt* posterior domain leads to the anterior shift of stripe 5 in *Kr* heterozygotes (Fig. 4G, blue). Meanwhile, the anterior boundary of stripe 2 remains comparable between heterozygotes and wild-types because of the unaltered anterior *gt* domains. While both stripes are directly regulated by *Kr*, this indirect effect through changes in *gt* expression contributes to a more significant effect on stripe 5 compared to stripe 2.

Collectively, our study demonstrates how decreased *Kr* dosage affects *eve* transcription dynamics through multifaceted interactions. While the direct regulation of Kr alters the positions of *eve* stripe 2 and 5 boundaries, indirect regulation via changes in other gap gene dynamics leads to a more dramatic effect on both the positions and transcription of *eve* stripes (Fig. 4G). Our results suggest that both the spatial and temporal patterning of early embryos are sensitive to fluctuations in transcription factor concentration, and the phenotypic consequences result from more convoluted interactions among multiple proteins and their intricate dynamics.

## Materials and methods

### Fly strains

Existing *eve-MS2* (Lim et al., 2018b), *gt>MS2-yellow* (*gt>MS2*) (Syed et al., 2021), *kni>MS2-yellow* (*kni>MS2*) (Syed et al., 2021), *nos>MCP-GFP, mCherry-PCP, His2Av-eBFP* (*MCP, PCP, His2Av*) (Lim et al., 2018a), Sp/CyO; Dr/TM3 (Bloomington Stock Center, #59967), and *Kr^1^/SM6* (Bloomington Stock Center, #3494) fly lines were used in this study. iab5>*snaPr-PP7-yellow* (*iab5>PP7*) reporter construct was generated by using the iab5 enhancer (Zhou et al., 1996), 100bp core snail promoter and *PP7-yellow* reporter gene (Keller et al., 2020). The plasmid was targeted to the VK02 locus in the second chromosome using PhiC-mediated site directed integration (Venken et al., 2006).

#### Imaging *eve-MS2* in wild-type and *Kr* heterozygous embryos

*iab5>PP7* virgin females were mated with Sp/CyO; Dr/TM3 males. *MCP, PCP, His2Av* virgin females were mated with Sp/CyO; Dr/TM3 males in parallel. The resulting *iab5>PP7/CyO; +/Dr* and *+/Sp; MCP, PCP, His2Av/TM3* offsprings generated from these two parallel crosses were then mated and homogenized to produce stable strain of *iab5>PP7*; *MCP, PCP, His2Av*. iab5>PP7; *MCP, PCP, His2Av* virgin females were mated with *Kr^1^/SM6* males to generate wild-type and *Kr* heterozygous embryos. The resulting *Kr^1^/iab5>PP7; +/MCP, PCP, His2Av* virgin female progenies were selected to mate with *eve-MS2* males. Embryos laid from this cross that inherit the *Kr^1^* allele are heterozygous for *Kr*, while embryos that inherit the wild-type allele are wild-type embryos and can be recognized by the *iab5>PP7* reporter. Both wild-type and *Kr* heterozygous embryos carry one copy of *eve-MS2* for visualization of *eve* expression.

#### Imaging gap genes in wildtype and *Kr* heterozygous embryos

*Kr^1^/iab5>PP7; +/MCP, PCP, His2Av* were generated using the same steps in the last section. The virgin females were collected and mated with *kni>MS2* or *gt>MS2* males. Embryos from this cross were used for imaging. Similarly, wild-type embryos were identified by the expression of *iab5>PP7*, while *Kr* heterozygous embryos have no *iab5>PP7* transcription. The embryos from crossing *Kr^1^/iab5>PP7; +/MCP, PCP, His2Av* with *gt>MS2* were also used in fluorescent *in situ* hybridization to detect *hb* and *Kr* mRNA.

### Live imaging

Embryos were collected 2 hours after being laid. They were then dechorionated with 50 % bleach for 2.75 minutes and mounted in Halocarbon oil 27 (Sigma) between a semipermeable membrane (Sarstedt) and coverslip (18 mm × 18 mm). All movies were imaged at room temperature (∼23 °C) with a Zeiss LSM800 confocal laser scanning microscope and Plan-Apochromat 40×/1.3 numerical aperture (N.A.) oil-immersion objective. 408 nm, 488 nm and 561 nm laser were used to visualize His2Av-eBFP2, MCP-GFP, and mCherry-PCP, respectively. To capture the full embryo, two adjacent tiles with 50 pixel overlaps were taken, resulting in a final image size of 950 x 500 pixels. A stack of 15 images with 0.7 µm step size in z were captured at each time point with a time resolution of 61s/ frame. For *eve-MS2*, seven biological replicates were taken for the wild-type embryos and five were taken for the *Kr* heterozygous embryos. Three replicates were taken for the wild-type and *Kr* heterozygous embryos for *kni>MS2* and *gt>MS2*. The same laser setting was used between wild-type and *Kr* heterozygous embryos.

### Fluorescent *in situ* hybridization

Embryos were collected 3 hours after being laid and fixated with formaldehyde. FISH was performed using previously published protocols (Lim et al., 2015). *Kr*-DIG, *hb*-FITC, *eve*-Biotin, *yellow-*Biotin RNA probes and Alexa Fluors were used to visualize *Kr*, *hb*, *eve,* and *yellow* expression. *yellow* probe was used to visualize *iab5>PP7-yellow*. Embryos were imaged with a Zeiss LSM800 confocal laser scanning microscope and Plan-Apochromat 20x/0.8 N.A. objective. Single image size was set to 1024 x 1024 pixels. Only the laterally oriented embryos in mid-NC14 were analyzed to minimize the noise due to orientation and developmental stage variations.

### Image analysis

Image processing and data analysis were performed in FIJI (ImageJ) and MATLAB (R2021b), using custom image analysis code (Keller et al., 2020). Images with maximum projections of Z-stacks were used for analysis.

#### Live imaging

Nuclei segmentation, nuclei tracking, and signal measurement were adapted from (Keller et al., 2020). MS2 signal within each nucleus was measured as the average of the two pixels with the highest fluorescent intensity from a max-projected image. To normalize for the variation in embryo size, each gene expression domain was measured as % egg lengths (EL). EL was measured as the distance between the anterior and posterior tips of each embryo. Each embryo was divided into 50 EL bins (2 %) along the AP axis. Gene expression profile along the AP axis was quantified as the average of the signal intensity of all nuclei within each bin at a given frame. Nuclei expressing active transcription in the last frame before gastrulation were used to define mature stripes. Each stripe domain is defined as the region between the anterior border of one mature stripe to the anterior border of the next mature stripe (e.g, the region between the anterior border of stripe 1 and the anterior border of stripe 2 is defined as “stripe 1 domain”). Stripe 7 region is defined as the anterior border of stripe 7 to the posterior end of the embryo.

Since each embryo has slightly different duration of NC14, the time from the end of the 13th mitosis to the beginning of gastrulation was normalized to 100 time points. Time after the onset of NC14 is labeled for snapshots of representative embryos (Fig. 1A-B, Fig. 2A-B, Fig. 3A). except for representative embryo images. mRNA production of individual nuclei was approximated by taking the integration of the area under the MS2 fluorescent intensity trajectory over time. mRNA production along the AP axis was measured as the average mRNA production of all nuclei within each bin.

At each time point, active nuclei were defined with an MS2 signal above a threshold value. Average transcriptional amplitudes were calculated as the mRNA output divided by the duration of actively transcribing time points. To track the boundary positions and width of each *eve* stripe over time, embryos were divided into five sections along the dorsoventral (DV) axis. AP axis positions of the most anteriorly and posteriorly located transcriptionally active nuclei were marked for each section. The average position of the five sections was used as a boundary position for each stripe.The stripe width was measured as the average distance between the anterior and posterior boundaries of each stripe. For Fig. 4 B-C, the boundaries for *kni* and *gt* posterior domains were set at the positions where the binned MS2 signal of *kni* or *gt* went above a threshold value.

#### Fluorescent *in situ* hybridization

The middle 40 pixels along the DV axis of each embryo image were used to measure *hb* and *Kr* RNA signal intensity. Embryos were divided into 1 % EL bins along the AP axis, and the signal intensities of all pixels within each bin were averaged. To minimize the effect of background noise, *hb* and *Kr* signals were subtracted by the lowest value from 10-90 % EL.

## Acknowledgements

We thank members of the Lim lab for helpful discussions and comments on the manuscript. We thank Hao Deng for preliminary studies. This study was supported by National Science Foundation CAREER MCP 2044613 awarded to B.L.

## Supplementary figure legends

**Fig. S1.**
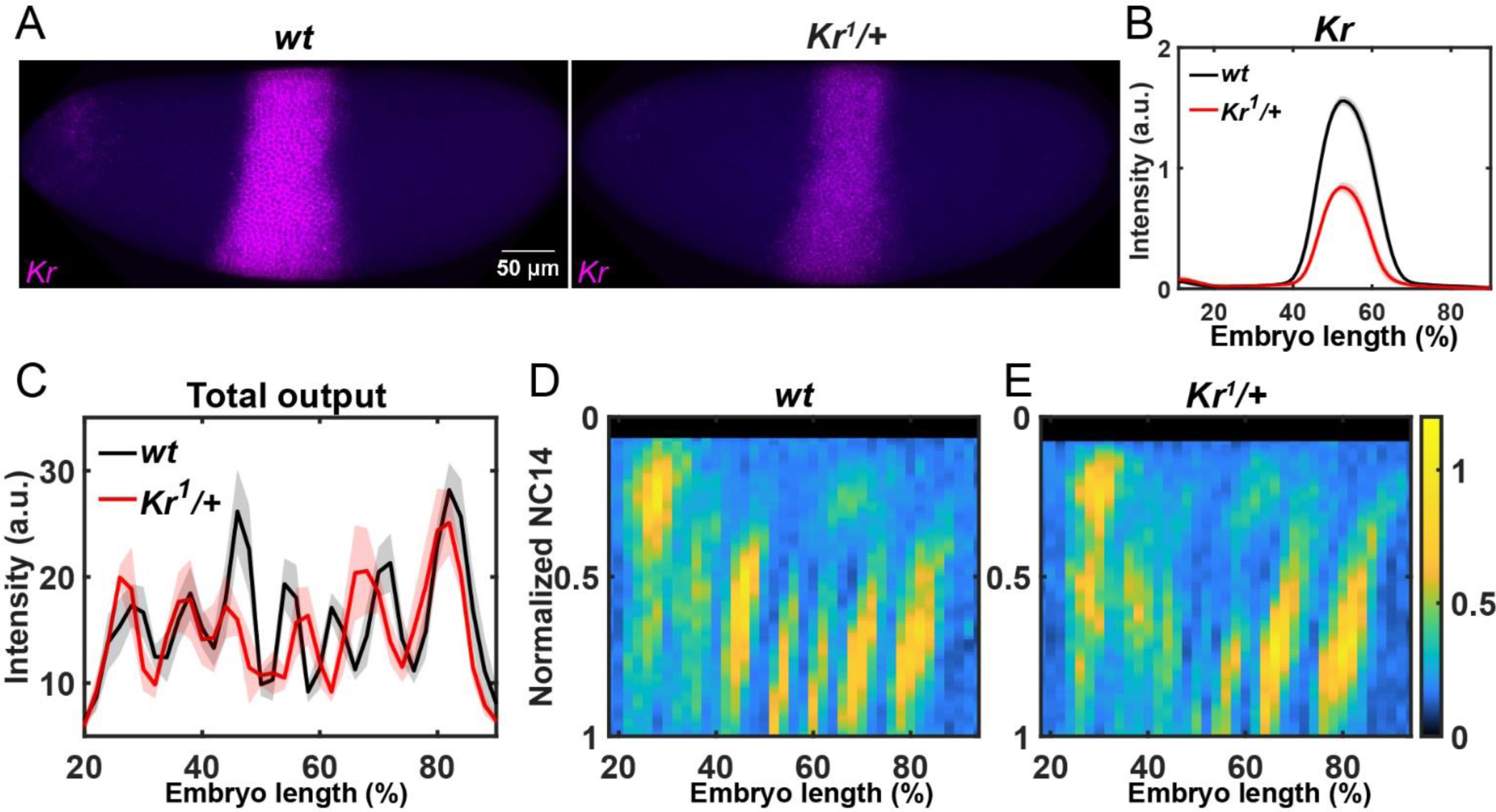
Decreased *Kr* dosage changes *eve* dynamics. (A) Mid-NC14 wild-type and *Kr* heterozygous embryo stained with *Kr* (magenta) and DAPI (blue). (B) Average spatial profile of *Kr* signal intensity. (C) Average spatial profile of *eve* mRNA production in the second half NC14. Shaded error bars show the mean ± s.e.m. of 10 wild-type and 7 *Kr* heterozygous embryos (B), 7 wild-types and 5 *Kr* heterozygous embryos (C), respectively. (D) Heatmaps showing the average spatial profile of eve-MS2 signal intensity in a wild-type and a *Kr* heterozygous embryo.

**Fig. S2.**
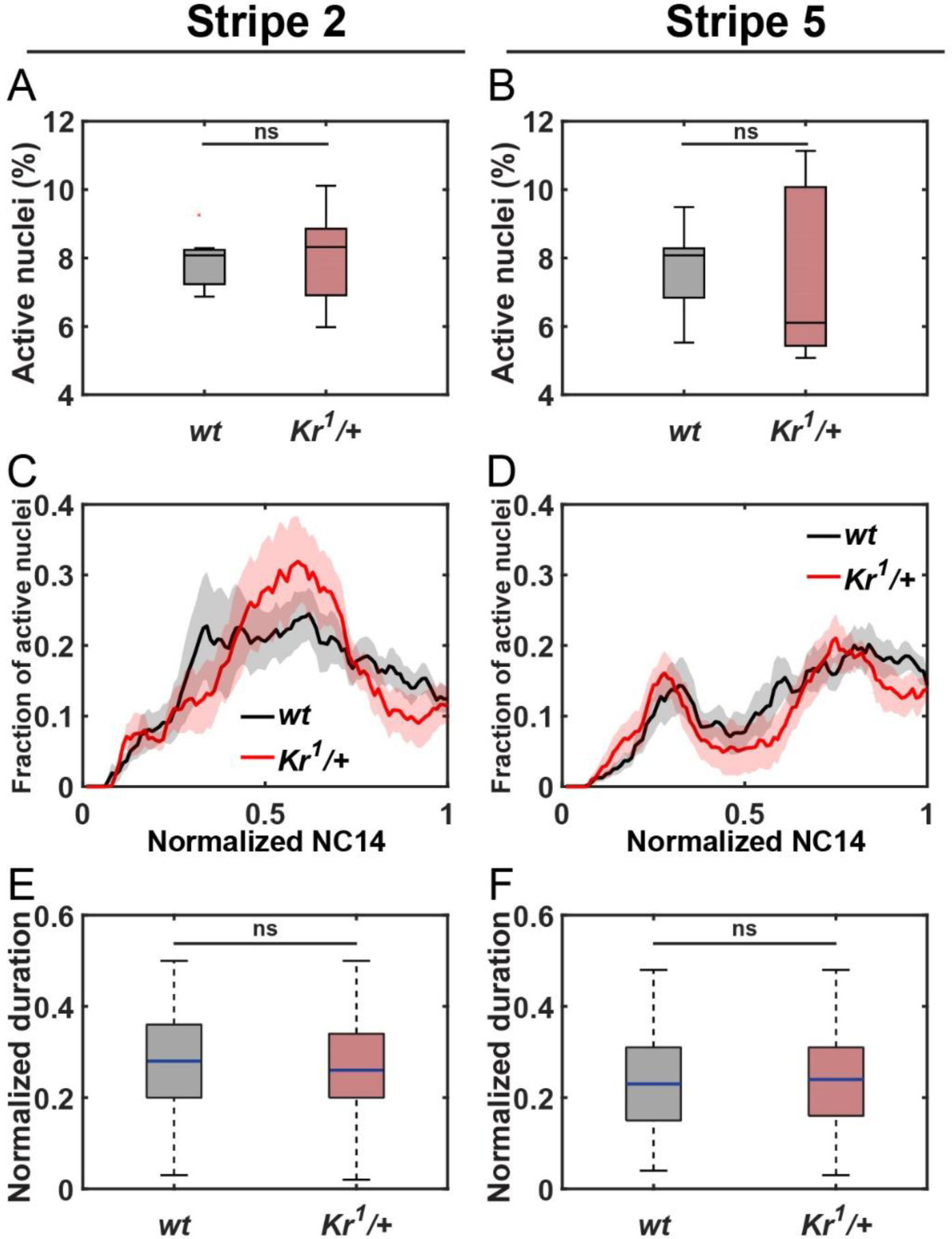
Decreased *Kr* dosage has limited effect on eve stripes 2 and 5 mRNA production. (A-B) Boxplots showing the percentage of transcriptionally active nuclei in stripe 2 (A) and stripe 5 (B), out of the total number of nuclei being analyzed in each embryo. (C-D) Average activation kinetics of stripe 2 (C) and stripe 5 (D). Shaded error bars represent mean ± s.e.m.. (E-F) Boxplots showing the transcriptionally active duration of individual nuclei in stripe 2 (E) and stripe 5 (F). ns, not significant from the Student’s t-test. The numbers of nuclei analyzed are 599 (stripe 2 *wt*), 498 (stripe 2 *Kr^1^/+*), 572 (stripe 5 *wt*), and 467 (stripe 5 *Kr^1^/+*). 7 wild-type and 5 *Kr* heterozygous embryos were analyzed.

**Fig. S3.**
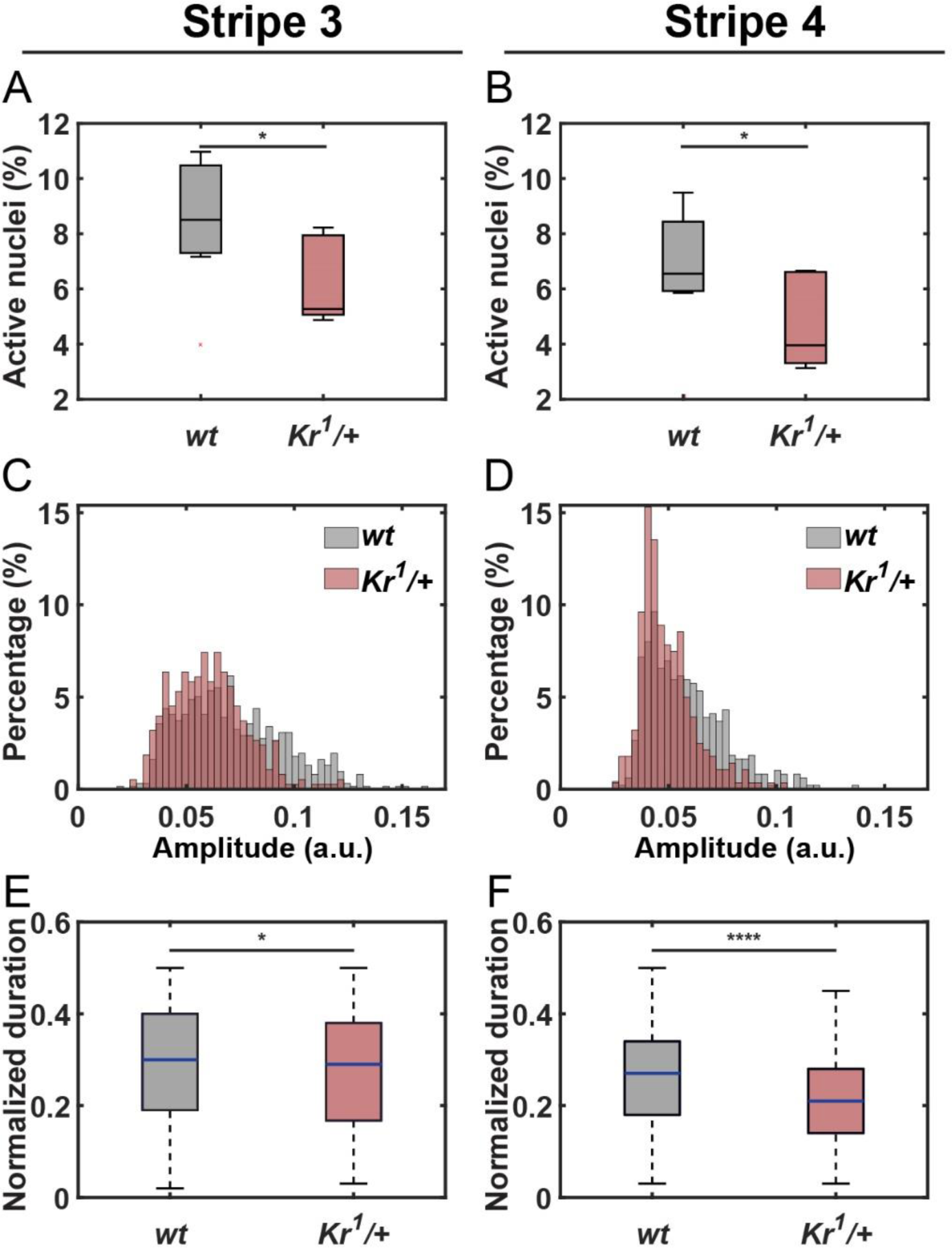
Decreased *Kr* dosage affects *eve* stripes 3 and 4 expression. (A-B) Boxplots showing the percentage of transcriptionally active nuclei in stripe 3 (A) and stripe 4 (B), out of the total number of nuclei being analyzed in each embryo. (C-D) Histograms showing the distribution of the average transcriptional amplitude of individual nuclei in stripe 3 (C) and stripe 4 (D). (E-F) Boxplots showing the transcriptionally active duration of individual nuclei in stripe 3 (E) and stripe 4 (F). *P<0.05 and ****P<0.0001 from the Student’s t-test. The numbers of nuclei analyzed are 617 (stripe 3 *wt*), 377 (stripe 3 *Kr^1^/+*), 488 (stripe 4 *wt*), and 281 (stripe 4 *Kr^1^/+*). 7 wild-type and 5 *Kr* heterozygous embryos were analyzed.

**Fig. S4.**
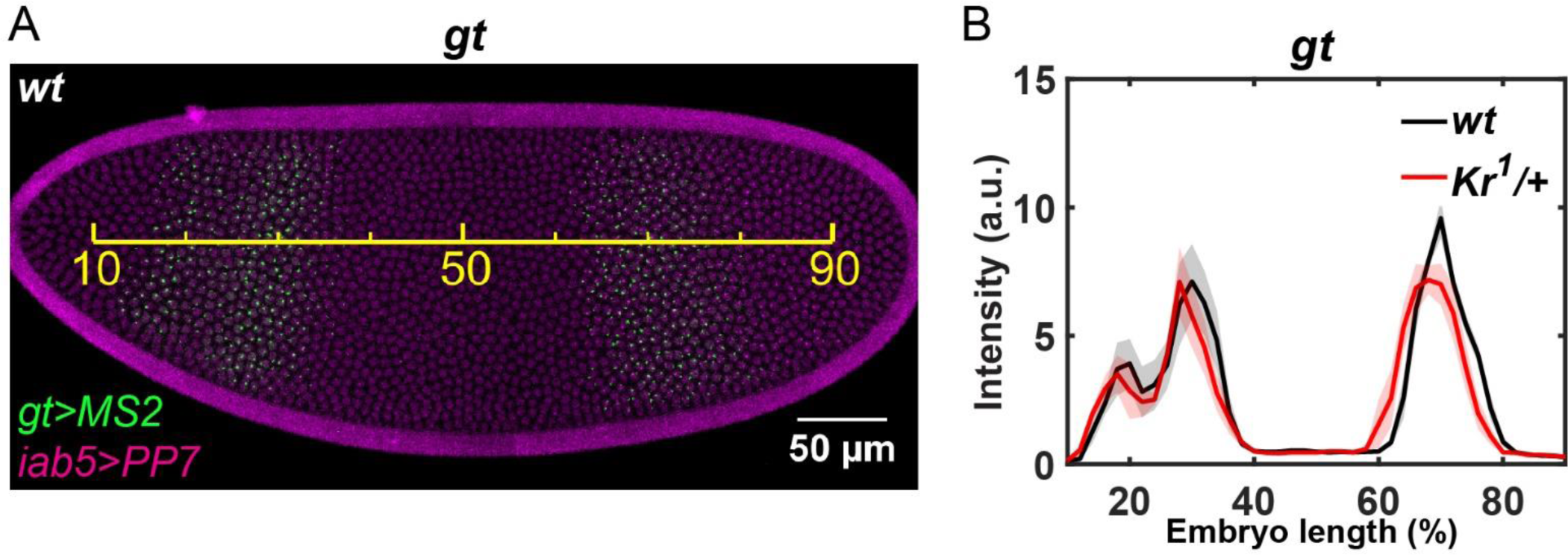
Decreased *Kr* dosage affects the posterior *gt* domain. (A) Snapshot of a wild-type embryo expressing *gt>MS2* (green) and *iab5>PP7* (magenta). Yellow scale bar represents the estimated EL across the AP axis. (B) Average spatial profile of *gt* signal intensity. Shaded error bars show the mean ± s.e.m.. 3 wild-type and 3 *Kr* heterozygous embryos were analyzed.

## Supplementary movie legends

**Movie S1.** Live imaging of *eve-MS2* in wild-type (top) and *Kr* heterozygous (bottom) embryos during NC14. *eve-MS2* signal is shown in green. *iab5>PP7* signal is shown in magenta. Embryos are oriented left-anterior, right-posterior.

**Movie S2.** Live imaging of wild-type (top) and *Kr* heterozygous (bottom) embryos during NC14. Transcriptionally active nuclei within the *eve* stripe 3 and 4 regions are false-colored in red, and other active nuclei are false-colored in gray. Embryos are oriented left-anterior, right-posterior.

